# Value-Based Neural Representations Predict Social Decision Preferences

**DOI:** 10.1101/2022.09.28.509596

**Authors:** João F. Guassi Moreira, Adriana S. Méndez Leal, Yael H. Waizman, Sarah M. Tashjian, Adriana Galván, Jennifer A. Silvers

**Affiliations:** Department of Psychology, University of California, Los Angeles; Department of Psychology, University of Southern California; Department of Humanities & Social Sciences, California Institute of Technology

**Keywords:** Social Decision-Making, Value, Pattern Expression, fMRI, Close Relationships

## Abstract

Social decision-making is omnipresent in everyday life, carrying the potential for both positive and negative consequences for the decision-maker and those closest to them. While evidence suggests that decision makers use value-based heuristics to guide choice behavior, very little is known about how decision makers’ representations of other agents influence social choice behavior. We used multivariate pattern expression analyses on fMRI data to understand how value-based processes shape neural representations of those affected by one’s social decisions and whether value-based encoding is associated with social decision preferences. We found that stronger value-based encoding of a given close other (e.g., parent) relative to a second close other (e.g., friend) was associated with a greater propensity to favor the former during subsequent social decision-making. These results are the first to our knowledge to explicitly show that value-based processes affect decision behavior via representations of close others.

---

Human beings are intrinsically social creatures and our decisions often have consequences for others. Such decisions—whose social consequences are direct or indirect—has been termed social decision-making. Neuroscientific research on social decision-making has increased dramatically over the past two decades, commensurate with its importance to both individual well-being (Lamba et al., 2020; Ong et al., 2017) and societal good (e.g., Johnson & Mislin, 2011). However, little research has examined the neural and behavioral underpinnings of decision making involving close others, instead focusing on decisions about strangers. This is surprising given that in everyday life we ostensibly care most about social choices that impact those closest to us. Moreover, despite the critical role that social and cognitive representations play in motivating social behavior, broadly construed (Guthrie et al., 2022; Tamir & Thornton, 2018), almost no research has examined how a decision-maker’s *representations* of others drives social decision processes. The present study sought to address these two gaps in the literature by testing how multivariate neural representations of two close others (a parent and a friend) predict subsequent social decisions about said others.

Most neuroscientific studies of social decision-making to date have paired behavioral paradigms from psychology and behavioral economics with neuroimaging to infer the underlying neural mechanisms supporting social decisions. One common approach involves fitting a computational model of decision-making to behavioral data, and subsequently examining the neural correlates of its parameters. A second, less common approach, is ‘model-agnostic’ and instead measures how brain activity is recruited during various task conditions. Both approaches have informed our understanding of social decision making – allowing us to observe seemingly fundamental rules such as preventing harm to others, obviating social uncertainty, and minimizing negative affect such as guilt or regret (Crockett et al., 2014; Feldmanhall & Chang, 2018; Lamba et al., 2020). Despite this, we argue these approaches have overlooked a critical component of social decision-making: how the decision-maker’s *representations* of other agents’ influences decision behaviors involving said agents.

Here, we define “representations” as internal models of others that dynamically integrate past and current information to guide prediction and future behavior (Clark & Toribio, 1994; DeCharms & Zador, 2000; Morgan, 2014; Poldrack, 2021). Representations have been shown to affect behavior in several adjacent fields such as cognitive science and social psychology. For instance, representations of similarity between another and oneself is thought to influence decisions about giving to others (Hackel et al., 2017). Similarly, subject-idiosyncratic representations of familiar, everyday objects predict between-object similarity judgments (Charest et al., 2014). Thus, this prior work from adjacent fields suggests how a decision maker internally represents specific individuals impacts social behaviors involving said individuals, though this assumption has rarely been explicitly tested.

It is thus likely that social decision preferences are also driven by representations in general, and value-based representations in particular. Social neuroscience research shows that brain regions involved in processing value such as the ventral striatum and medial prefrontal cortex are critical for encoding and tracking social information about other individuals (Zerubavel et al., 2015) as well as supporting cognitive heuristics during social decision-making (Chang et al., 2011; Fareri et al., 2015). Given the importance of value-based computations in social behavior, and the significance of representations in driving decision making, it seems likely that value-based representations play a key role in coordinating social decision making behavior. Specifically, the literature begs the question of whether stronger value-based encoding of representations of specific others (e.g., parents, friends) is linked to social decision preferences for said others. However, this possibility has yet to be formally tested.

In the current study, we sought to determine the extent to which neural representations of two specific close others (parents and friends) were encoded as neural signatures of valuation, and related these estimates to social decision-making preferences involving these others. Specifically, we examined whether value-based representations of parents and friends predicted whether individuals would prioritize one close other at the expense of another. We elected to focus on this type of decision scenario (pitting a parent versus friend) for two reasons. First, the vast majority of social decision research to date has focused on social decisions about unfamiliar others, rather than close others. However, recent studies indicate that social decision behavior often changes as a function of whom is affected (Fareri et al., 2020, 2022; Powers et al., 2017; van de Groep et al., 2022), As such, we chose to focus on close others so as to increase the generalizability of social decision research. Second, we have conducted extensive behavioral work specifically examining social decisions between parents and friends (Guassi Moreira et al., 2018, 2020, 2021), having previously observed a general tendency among older adolescents to favor a parent over a friend, in addition to considerable heterogeneity in the direction and magnitude of individual preferences. That this particular decision scenario is well studied—and has been replicated—renders it an ideal test case for the current study. We hypothesized (pre-registered; osf.io/muv2c) (1) that individual neural representations of parents—relative to friends— would be more strongly expressed as neural signatures of value, and (2) that individual differences in value-based expression in neural representations would predict social decision preferences (i.e., greater value-based expression in one’s neural representation of a close other will be associated with a greater tendency to favor said close other). Testing these two hypotheses stands to enrich our understanding of how value-based representations drive consequential social decision making behavior, especially in contexts that have relatively greater ecological validity (e.g., navigating decisions with conflicting outcomes for multiple close others).

## Methods

### Overview

The goal of this study was to examine the role of value-based neural representations in social decision-making preferences. We did this using pattern expression analyses (Doré et al., 2017; Hong et al., 2019; Cosme et al., 2020). Pattern expression analyses are commonly used to answer questions about how strongly a given brain state is expressed as a psychological process of interest, resulting in a single score that is used to compare relative differences in expression between brain states (Cosme et al., 2019; Doré et al., 2017). For this study, our intent was (i) to determine how strongly neural representations of parents and friends were expressed as signatures of value and then (ii) examine whether the scores could predict social decision preferences.

### Participants

Participants for this study were comprised of 48 older adolescents (18-19 years). We targeted older adolescents because theoretically heightened social sensitivity to peer processes coinciding with continued reliance on parental relationships makes this developmental stage an ideal phase during which to examine social decision behavior that pits the interests of two relatively important close others (Blakemore & Mills, 2014; Steinberg & Morris, 2001). Participants were recruited by posting flyers and sending mass emails to undergraduate college students. In order to be eligible to participate, individuals were required to (i) be between the ages of 18 and 19 years old, (ii) be eligible for MRI scanning (e.g., no metal implants, no claustrophobia, etc.), (iii) be a fluent English speaker, (iv) have no neurological impairments, (v) be able to nominate two close others (a parent and friend) and provide photographs and names for each (more information about the nomination procedure and stimuli follow below). Participants were compensated with a $25 (USD) cash payment plus an additional $1-5 bonus chosen at random (described in greater detail below). Three participants were excluded from all analyses (one because of a scanner computer error, a second due to poor overall data quality, and a third due to discovery of a biological artifact), resulting in a final sample size of 45. All participants provided written consent in accordance with the policies of the UCLA Institutional Review Board.

### Sample Size Considerations

The best practices for determining sample size in human neuroimaging research are relatively unclear given the complexity and difficulty in calculating power for task-based fMRI studies (Chen et al., 2017; Cremers et al., 2017; Mumford, 2012; Poldrack et al., 2017). Recent research suggests very large sample sizes are needed to examine individual differences between resting state fMRI and behavior (Marek, Tervo-Clemmens, et al., 2022), yet it is unclear how this finding generalizes to task-based analyses, multivoxel fMRI approaches, or analyses that involves repeated behavioral measurements nested within subjects. Further complicating matters is the fact that fMRI is a particularly expensive neuroimaging modality. Given these realities and the lack of clear sample size requires, our goal set prior to data collection was to scan as many participants as our funding would allow, preferably exceeding the most recently estimated median cell size in human fMRI research (*N* = 35; Poldrack et al., 2017). We acquired funding to scan 50 participants, but stopped data collection early in light of the COVID-19 pandemic.

### Experimental Protocol

#### Overview

Participants were asked to nominate a parent and close friend of their choice, and provide stimuli (photos, names) of each person prior to their scheduled scan date. Participants completed a fMRI task to elicit neural representations of their parent and friend (Parent-Friend Representation Task, described below), as well as another fMRI task which was used to define a sample-specific neural signature of value (Coin Flip Task, also described below). Last, participants completed a post-scan session to assess behavioral social decision preferences. Each element of this procedure is described in greater detail below.

#### Parent-Friend Nomination and Stimuli Collection

Participants were instructed to nominate a parent and a close friend, and provide custom stimuli of each of them for an fMRI task. Details about these nominations and the stimuli can be accessed in the Supplemental Information.

### fMRI Tasks

#### Parent-Friend Representation Task

In a block design, participants were shown custom stimuli of their own parent and nominated friend to elicit and record neural representations of each close other. In a given block, participants saw randomly ordered stimuli pertaining to one close other (parent or friend). Various elements of this task were designed to be broadly consistent with prior social and affective neuroscience literature (Gee et al., 2014; Parkinson et al., 2017; Taylor et al., 2009; Zerubavel et al., 2015). These stimuli were comprised of the five headshots in addition to the close other’s name^1^ printed in five unique fonts – ‘Berlin’, ‘Broadway’, ‘Calibri’, ‘Colonna’, and ‘Comic Sans’ (10 unique stimuli). The use of varying photographic and text stimuli was intended to elicit amodal neural representations of parents and friends, thereby avoiding basic perceptual confounds. Each block contained 20 rapid presentations (2 for each of the 10 unique stimuli) of said stimuli (1000ms) with a brief inter-stimulus interval (ISI) between images (500ms). Participants completed a one-back task based on stimulus type (photo vs text, regardless of orientation or font) to ensure they were paying attention (i.e., press a button if the current stimulus type matches the one shown just before it). 15000ms of fixation between blocks was presented to account for lagged effects of the hemodynamic response function. Six blocks (3 parent, 3 friend) and six inter-block fixation periods were presented per run. As a result, the entire task lasted approximately 4.5 minutes (270s): [1500ms/trial x 20 trials/block x 6 blocks] + [15000ms inter-block fixation periods x 6 fixation periods].

#### Coin Flip Task

Following the representation task, participants completed two runs of a reward task intended to evoke neural representations of value (Braams & Crone, 2016). During this event-related task, participants guessed the outcome (‘Heads’ or ‘Tails’) of a series of coin flip gambles in order to win or lose monetary rewards (presented as coins). Each trial began with a reward summary (3000ms), a screen that lists the amount awarded or lost for guessing correctly or incorrectly, respectively. Participants made their guess, via button press, at this stage (‘Heads’ or ‘Tails’). Following a 1000ms inter-stimulus interval, participants received feedback about whether their guess was correct or incorrect (2500ms). A jittered inter-trial interval separated trials, with values drawn from an exponential distribution (mean = 2880ms, SD = 2660ms, range = 1000-10000ms). Each run lasted approximately 6 minutes. Participants completed 30 trials per run, broken down across three distinct trial types: (i) win 3 coins, lose 3 coins; (ii) win 5 coins, lose 2 coins; (iii) win 2 coins, lose 5 coins. Participants were told the coin is fair (i.e., P(‘Heads’) = ½). In reality, the task was rigged such that individuals won approximately half of the trials to ensure enough gain and loss events for subsequent modeling and estimation. To obtain a generalized signature of valuation, one run varied the type of coins (Kennedy coin vs Sacagawea coin) and thus the perceptual features of the coin (color: silver vs gold; gender of the head: male vs female; etc.). The orientation of the coin also varied for this reason (i.e., half of the reward summaries showed the coins on the ‘Heads’ side, the other half showed them on the ‘Tails’ side). Last, participants were informed a subset of the trials would be selected at random and added to, or subtracted from, their earnings (up to +/- $5). In actuality, participants always received a randomly selected bonus between $1 - $5.

#### fMRI Data Acquisition

Neuroimaging data were collected using a research-dedicated 3 Tesla, Siemens Magnetom Prisma MRI scanner and 32-channel head coil. A high resolution T1* magnetization-prepared rapid-acquisition gradient echo structural image was acquired for registration purposes (MPRAGE; TR = 2400ms, TE = 2.22ms, Flip Angle = 8º, FOV = 256 mm^2^, 0.8 mm^3^ isotropic voxels, 208 slices, A >> P phase encoding). Functional runs were comprised of T2*-weighted multiband echoplanar images (TR = 1000ms, TE = 37ms, Flip Angle = 60º, FOV = 208 mm^2^, 2.0 mm^3^ isotropic voxels, 60 slices, A >> P phase encoding, multi-band acceleration factor = 6). These parameters were informed by studies on related topics using similar analytic techniques (e.g., Chang et al., 2015; Chavez et al., 2017).

### Post Scan Procedure

Participants completed the following procedure directly after the fMRI scan. This procedure was intended to measure social decision preferences between the participants’ nominated parent and friend, and acquire additional information about these nominees.

#### Parent-Friend Salience Procedure

Before completing the social decision-making paradigm described below, participants answered brief prompts about the parent and friend they had nominated. This procedure was enacted to amplify the salience of completing the subsequent social decision-making task in the absence of their parent and friend, consistent with prior studies (Guassi Moreira et al., 2018, 2020). Participants provided basic information about each close others (e.g., name, age, sex), briefly wrote about a memory (∼1 paragraph) they share with each close other, and listed a handful of words and phrases describing each close other.

#### Social Decision-Making Paradigm

After scanning, and consistent with our prior behavioral work (Guassi Moreira et al., 2018, 2020), we used a modified version of the computerized “hot” Columbia Card Task (CCT) to assess social decision-making preferences involving conflicting outcomes for parents and friends (Figner et al., 2009; van Duijvenvoorde et al., 2015). The modified CCT is an iterative risk-taking task in which individuals turn over cards that can result in hypothetical rewards or losses. The modification we previously introduced applied a trade-off such that rewards exclusively benefitted one of the two close others and losses were exclusively incurred by the second of the two close others.

Participants in our study completed two runs of this task, one in which the rewards benefitted the parent and the losses were incurred by the friend, and a second run where the opposite was true (condition order counterbalanced between subjects). Because there is always a trade-off in the conflicting outcomes for the two close others, the task can be modeled to reveal whether there is an aggregate preference for a parent or friend. Technical details of the task and its administration can be accessed in the Supplemental Information. This was administered after the scan, with an experimenter present to unobtrusively monitor the participant.

#### Additional Measures

Participants completed a series of self-report measures on a laboratory computer via Qualtrics (an online survey administration platform), including measures of subjective relationship quality, domain-specific risk-taking for oneself, sensation seeking, and family obligation. Participants also completed a computerized risk-taking task that affected only themselves (i.e., self-oriented risks).

## Analysis Plan

### fMRI Data Preprocessing

Prior to preprocessing, data were visually inspected for artifacts and anatomical abnormalities. Data were preprocessed and analyzed using the fMRI Expert Analysis Tool (FEAT, Version 6.00) of the MFRIB Software Library package (FSL, Version 5.0.9; fsl.fmrib.ox.ac.uk). Preprocessing began by using the brain extraction tool (BET) to remove nonbrain tissue from functional and structural images, followed by head motion correction via spatial realignment of functional volumes using MCFLIRT. The data were hi-pass filtered to remove low frequency artifacts (45s for the Parent and Friend Representation Task; 100s for the Coin Flip task). From there, the extent of head motion artifacts was estimated by using the FSL Motion Outliers command to document volumes that exceed a 0.9 mm threshold of framewise displacement (FD; Siegel et al., 2014). Runs with 25% of volumes exceeding this threshold were excluded from analysis. Head motion in the sample was low overall: the ‘average subject’ moved less than one volume above the threshold with a maximum FD value of 0.6 (full descriptive information about head motion can be accessed in the Supplemental Information). To help reduce high frequency noise introduced by realignment (Etzel et al., 2011; Misaki et al., 2014), data were smoothed with a 1 mm Gaussian kernel (full width at half maximum). Data were pre-whitened prior to analysis to correct for autocorrelated residuals. FSL’s boundary based registration algorithm (Greve & Fischl, 2009) was used to register functional data to the high resolution structural scan (MPRAGE). MPRAGE images were then nonlinearly registered to the MNI152 template image (10-mm warp resolution), and the ensuing transformation matrix was used to register functional images to standard space. This step also resampled voxel size to 2mm^3^ isotropic.

All participants had usable data for the Parent and Friend Representation Task, although three participants only had 1, 2 and 3 usable runs (out of four), respectively, of the task available for analysis. Three participants were excluded from analyses involving the Coin Flip task. Two such participants were excluded because they lowered part of their heads out of the coil during the Coin Flip task, rendering missing data for large parts of the temporal pole. The third such participant was excluded due to head motion, as they averaged 22 volumes exceeding the FD threshold (average maximum FD = 8.82mm) across both runs. The final sample size for all analyses was *N* = 45.

### Multivariate Pattern Estimation

We used our data to estimate three multivariate neural patterns: a parent representation, a friend representation, and a value-based signature.

#### Parent and friend representations

Estimating the parent and friend neural representations was accomplished by modeling the Parent and Friend Representation Task with a standard General Linear Model (GLM) analysis. Each run of the task was submitted to a fixed effects GLM analysis in FSL. Parent and friend blocks were modeled with respective boxcar regressors, convolved with the hemodynamic response function (double gamma) and bandpass filtered to avoid reintroducing noise into the data. Slice timing effects were addressed by also modeling the temporal derivative of each task regressor. Head motion was statistically adjusted for by adding rotation and translation parameters, along with their derivatives and squares (obtained from MCFLIRT motion correction) as nuisance regressors. To further statistically adjust for potential spurious effects of head motion, we included additional regressors for individual volumes that exceeded the 0.9 mm FD threshold. Two linear contrasts were computed: parent > baseline and friend > baseline. A second level (subject level) analysis was carried out to average contrast estimates over the four runs, using a fixed effects model and forcing random effects variance to zero. The ensuing parent > baseline and friend > baseline maps, one each per subject, served as the estimates of parent and friend representations.

#### Value-based signature

We created a neural signature of value consistent with methods previously employed with similar tasks (Chang et al., 2015; Cosme et al., 2019; Wager et al., 2013; Reddan et al., 2018). This process involved training a statistical model to predict gain and loss values on each trial of the Coin Flip Task based on brain activity, and ultimately yielded a statistical map containing voxel weights that represent the strength of association between voxel activity and reward/loss outcomes.

The first step in this task was to compute brain activity for individual trials on the Coin Flip task. We accomplished this by conducting a least squares single (LSS) analysis (Mumford et al., 2012, 2014). Briefly, LSS entails creating a unique fixed effect GLM for every trial, in every run, for all participants^2^. We created a single-event regressor for a given trial in its respective GLM and model all other trials in their respective conditions. For the coinflip task, this meant that any given LSS GLM would contain a regressor for the current ‘target trial’, a regressor for gain outcomes, a regressor for loss outcomes, and a regressor for guessing between ‘Heads’ or ‘Tails’ (i.e., the length of presentation time for the reward summary). A linear contrast comparing trial > baseline was estimated for each GLM. The ensuing single-trial estimates from all participants were used to extract a *t* x *v* matrix, containing brain activity during the *t-*th trial in the *v*-th voxel (whole brain). Given the high dimensionality of this matrix (209,036 voxels), principal components analysis (PCA) was employed to reduce the number of features (i.e., voxels). Finally, penalized regression (e.g., LASSO, ridge) models were fit to the data, predicting the monetary outcome of each trial from its brain activity and thus yielding a set of weights for each principal component. Weights for each component were backtransformed into voxel space, yielding the final neural signature of value. Given some sparsity observed among the voxel weights, we created two additional versions of the map by smoothing the maps using a 2mm and 4mm Gaussian kernel (fwhm). All three versions are used in analyses reported below. More details about the signature creation process can be accessed in the Supplemental Information, along with a visualization of the unsmoothed signature (Figure 2).

**Figure 1.** Schematic of the two fMRI tasks. FIGURE OMITTED DUE TO BIORXIV POLICY OVER IDENTIFIABLE INFORMATION *Note*. ‘Representation Task’ refers to the Parent and Friend Representation task. The Representation Task was always administered before the Coin Flip Task.

**Figure 2.**
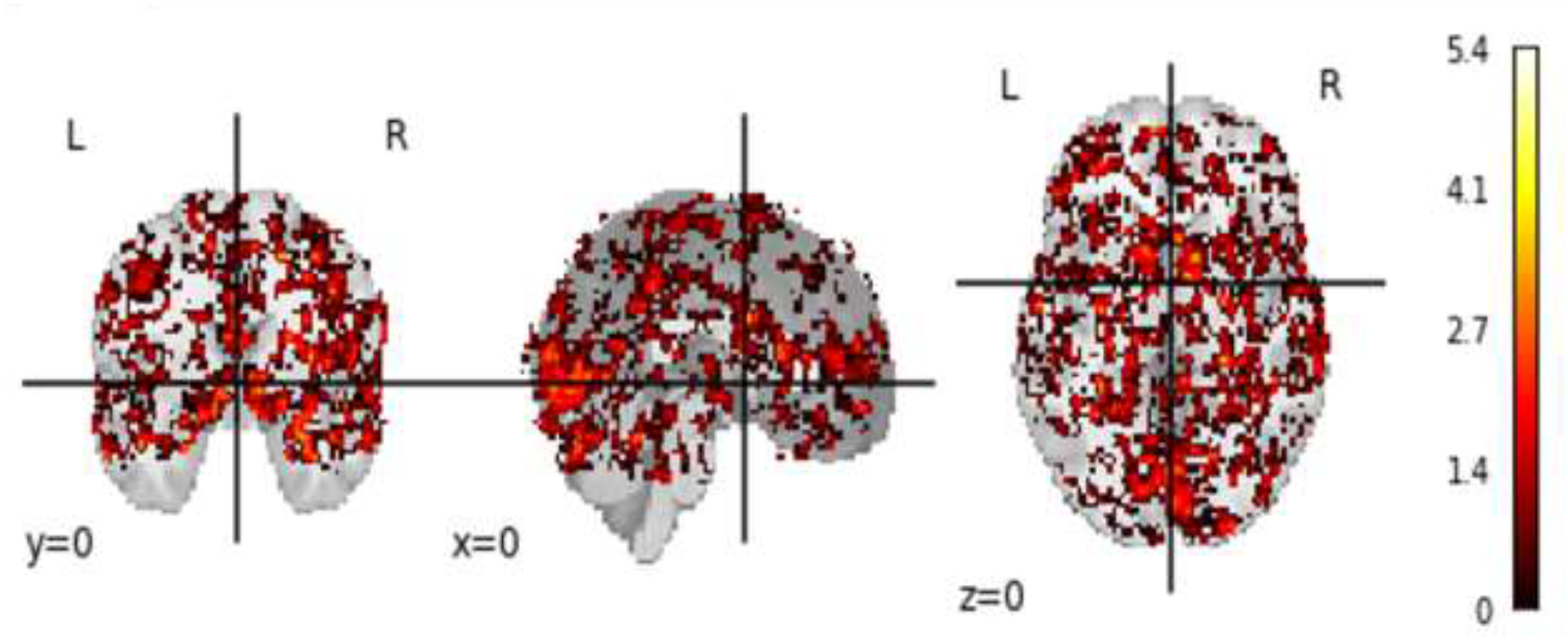
Sample-specific neural signature of value (unsmoothed).

#### Robustness checks

Two types of robustness checks were performed on our value signature methods. First, to ensure the signature was specific to value and did not inadvertently tap another psychological process, we cross-referenced its similarity with publicly available meta-analytic maps of similar and distinct constructs (see Supplemental Information). Second, we re-ran all analyses using two additional neural signatures of value, defined by Neurosynth (Yarkoni et al., 2011) to ensure our findings were robust to the method of signature definition (see Supplemental Information). Notably, we see merit in using a two-pronged approach to capturing neural signatures in that one signature is representative of the population of interest here (sample-specific) and another is based on data derived from thousands of participants (Neurosynth). As described below, results were largely consistent between these two approaches.

#### Pattern Expression Analysis

Pattern expression analysis captures how much a given psychological process (indexed by a neural signature) contributes to a representation or state. The analysis involves taking the voxel-wise dot product between values in a neural representation and a neural signature of interest. The computation is given by the following equation.

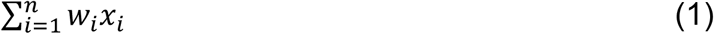

Where *n* is the number of voxels, *w*_*i*_ are the weights of the neural signature (value patterns created from the coin flip task data), and *x*_*i*_ is the neural activity (inferred via BOLD) from the representation’s voxels (for parent and friend representations, respectively).

#### Statistical Analysis

After extracting pattern expression scores, we first examined whether parent representations were more strongly encoded as signatures of value, relative to friend representations. We tested this by analyzing paired differences in parent – friend pattern expression scores.

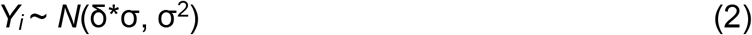

Here *Y*_*i*_ represents the paired pattern expression difference score for the *i*-th participant, and it is modeled as being drawn from a normal distribution, centered around a mean (δ*σ) and variance (σ^2^). The mean was parameterized as δ*σ so that draws from this distribution are in *Y*_*i*_’s ‘native units’, but the resulting summary statistics reflect standardized effect sizes (i.e., mean/standard deviation). The model assigned priors for both δ and σ. The variance was given a Jeffreys prior (p(σ^2^) ∝ 1/σ^2^), and δ— the mean effect size—was modeled as being distributed Cauchy (δ ∼ Cauchy(0, r), where r = 1/sqrt(2)). This model was fit using *rstan*, a package in the R statistical software library that allows the user to interface with *Stan* a Bayesian modeling software (Stan Development Team, 2020) (no thinning, 4 chains, 2,000 samples per chain, 1,000 discarded burn-in samples).

The next analytic step tested whether individual differences in pattern expression scores predicted social decision preferences. To this end, we used a hierarchical Bayesian model. Details of the model, including selection of priors, are described in the Supplemental Information.

#### Inference Criterion

Inference was performed on the posterior samples by using the region of practical equivalence method popularized by Kruschke (2011, 2013). We employed this method in three steps. First, a credible interval (CI)—a span of the posterior distribution capturing a user-defined portion of its mass—was computed for a given posterior using the Highest Density Interval (HDI) method (*bayestestR* package; Makowski et al., 2019). We used 89% credible intervals upon the recommendation that wider intervals (e.g., 95%) are more to sensitive Monte Carlo sampling error (Makowski et al., 2019; McElreath, 2018). Second, we specified a region of practical equivalence (ROPE), which is a user-defined interval in the parameter space whose values are deemed virtually equivalent to a null value (i.e., spans effects of such little magnitude that they are, for practical purposes, considered comparable to the null value). We defined a ROPE of [-0.1, 0.1] for analyses involved paired comparisons because we were uninterested in standardized effects below 0.1 in magnitude and a ROPE of [-0.095, 0.095] was defined for hierarchical logistic regression (i.e., a 10% expected change in the likelihood of flipping over a card after transforming logistic regression coefficients back into the odds scale). Finally, we inspected the degree of overlap between the CI and ROPE and compared it to the inferential criteria specified by Spiegelhalter and colleagues (1994). Here, if the CI falls completely outside of the ROPE, evidence for an effect is said to be robust (Kruschke, 2011), whereas if the CI overlaps with ROPE on side, then there is evidence to rule out parameter values only on the non-overlapping side of the ROPE. If the ROPE entirely contains the CI, then that is evidence in favor of accepting a null effect, and if the CI spans the ROPE but extends outside both ends of it, then the evidence is ‘equivocal’.

## Results

### Manipulation Checks

We conducted two key manipulation checks prior to executing the aforementioned analysis plan. First, we computed linear contrasts (win > loss) from a traditional univariate analysis of the Coin Flip task to ensure the task was recruiting brain regions previously implicated in valuation (Haber & Knutson, 2009; Knutson et al., 2001). Second, we analyzed behavioral data from the modified CCT data (collected post-scan) without any between-person predictors to check whether we could replicate a previously observed overall parent-over-friend preference (Guassi Moreira et al., 2018, 2020). Both manipulation checks suggested replication of prior findings, with imaging results showing robust activity in the ventral striatum and medial prefrontal cortex, and parameters of the behavioral decision-making model suggesting a parent-over-friend preference (see Supplemental Information). These results suggest any potential null effects in other analyses would not be due to the current sample exhibiting differing social decision preferences than those of samples that inspired the current study.

### Paired Differences in Value-Based Pattern Expression of Neural Representations

Using the five different neural signatures of value (3 sample-specific signatures created using different levels of smoothing; uniformity and association Neurosynth maps), we observed mixed evidence for the hypothesis that parent and friend neural representations are differentially encoded as a function of value, with more evidence in favor of friends than parents. Results using the sample-specific and Neurosynth neural signatures of value both showed a bias towards *friends*, not parents, as indicated by the mean of posterior samples.

The results were relatively stronger in favor of friends over parents with the Neurosynth signature than the sample-specific signature. Results using the Neurosynth maps as neural signatures showed that the majority of the posterior mass either fell within ROPE or in the negatively signed region encoding friend > parent relatively stronger evidence for a value-based bias in friend neural representations (*NS:* posterior mean, (SD): *d* = -0.24 (0.15), 89% CI: [-0.48, -0.01]) (*NS_Asc:* posterior mean, (SD): *d* = -0.14 (0.14), 89% CI: [-0.37, 0.09]). This was contrary to hypotheses, in that it suggested that *friend* representations are more strongly encoded as value-based signatures, Figure 3, bottom row).

**Figure 3.**
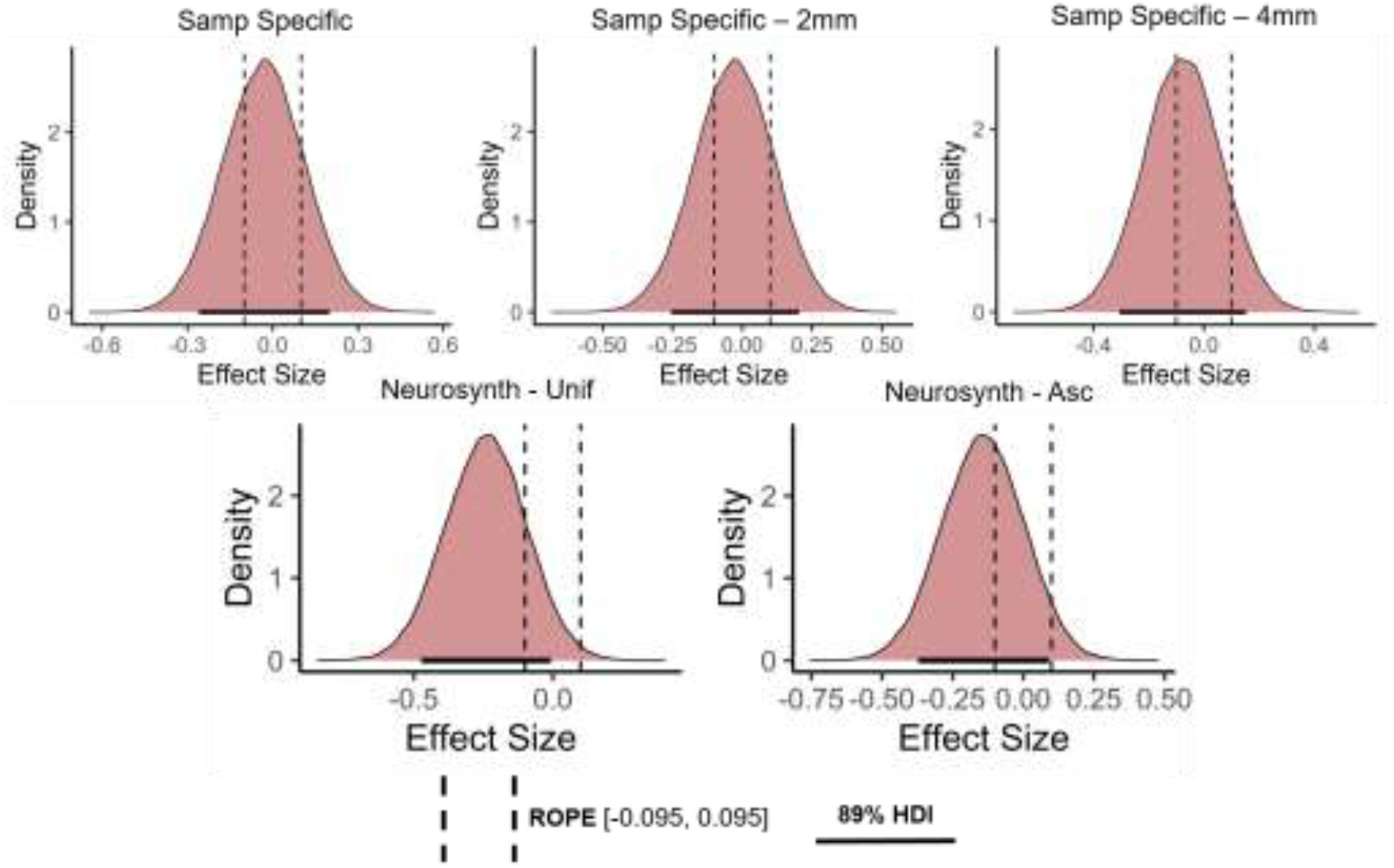
Posterior distributions of paired differences in value-based pattern expression values *Note*. ‘Samp Specific’ and ‘Neurosynth’ refer to the type of signature used (Samp specific = sample-specific signature built using ridge regression; NS = Neurosynth signature obtained from large scale, automated meta analysis). ‘2mm’ and ‘4mm’ refer to the degree of smoothing applied to the sample specific signature (the top left signature had no smoothing applied). ‘Asc’ refers to the Neurosynth association map; ‘unif’ refers to the Neurosynth uniformity map. Paired differences are in a standardized metric (*d*). ‘ROPE’ refers to Region of Practical Equivalence; ‘HDI’ refers to highest density credible intervals. Difference scores were computed by subtracting friend from parent (parent – friend).

For the sample-specific results, roughly equal amounts of the posterior mass lay on either side of ROPE, suggesting the evidence for an effect in either direction was equivocal (Figure 3, top row; *Samp Specific*: posterior mean, (SD): *d* = -0.08 (0.14), 89% CI: [-0.31, 0.15]) (*Samp Specific – 2mm*: posterior mean, (SD): *d* = -0.03 (0.14), 89% CI: [-0.25, 0.20]) (*Samp Specific – 4mm:* posterior mean, (SD): *d* = -0.03 (0.14), 89% CI: [-0.27, 0.19]). Again, these results were contrary to hypotheses. We conducted two *post-hoc* follow-up analyses to determine whether these unanticipated results could have been driven by brain regions in the neural signature that were potentially capturing a non-relevant psychological process, thereby obscuring relevant signal. However, neither post-hoc analysis substantially changed results (see Supplemental Information).

### Modeling Social Decision Preferences as a Function of Value-Based Pattern Expression

Pattern expression values derived using the three versions of the sample-specific neural value signature and the two versions of the Neurosynth neural value signature predicted subsequent social decision preferences on the modified CCT (Tables 1-2, Figures 4-5). Across multiple signatures, we observed that greater value-based pattern representation of a given close other predicted favoring said other in the modified CCT (e.g., greater value-based pattern expression for parent predicted a subsequent behavioral preference for parent on the CCT). For the sample-specific signatures, these results were observed on the two models using a smoothed neural signature value, whereas contradictory results were observed with an unsmoothed neural value signature. Based on our inferential criteria, greater parent value-based pattern expression was either related to equivocal preferences or parent preferences, but not friend preferences. Confidence is strengthened by the fact that a plurality of the posterior mass fell in the direction of the hypothesized effect (parent PE predicting parent preference) for all parameter estimates.

**Table 1.**
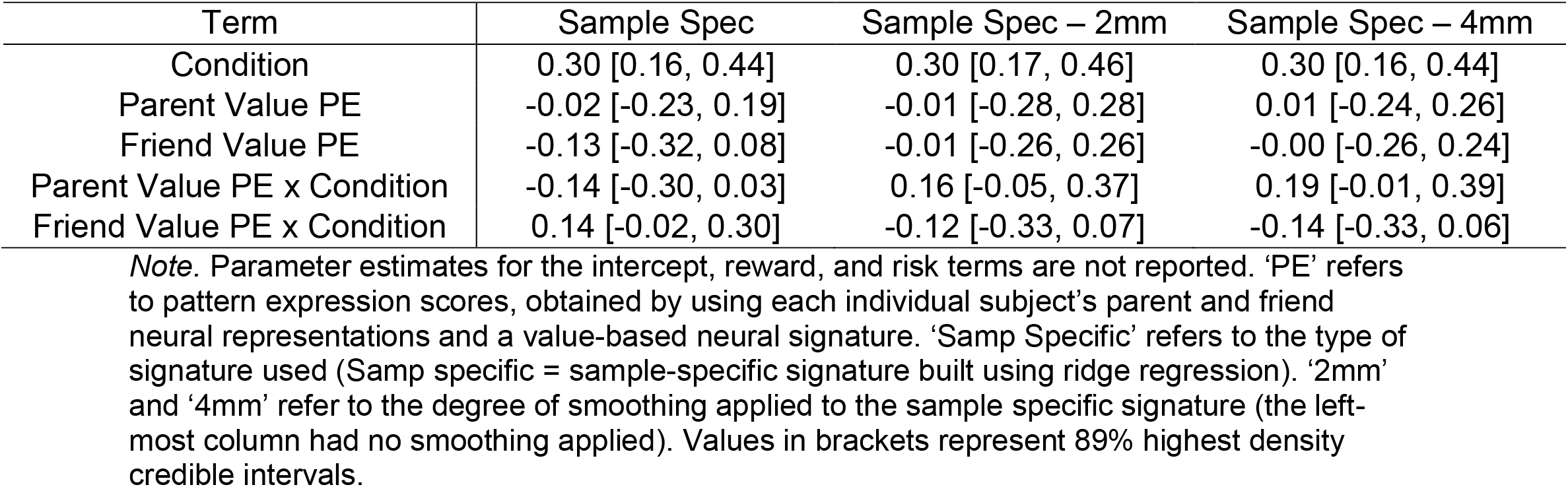
Predicting social decision preferences as a function of value-based representations using a sample-specific neural signature.

| Term | Sample Spec | Sample Spec – 2mm | Sample Spec – 4mm |
| --- | --- | --- | --- |
| Condition | 0.30 [0.16, 0.44] | 0.30 [0.17, 0.46] | 0.30 [0.16, 0.44] |
| Parent Value PE | -0.02 [-0.23, 0.19] | -0.01 [-0.28, 0.28] | 0.01 [-0.24, 0.26] |
| Friend Value PE | -0.13 [-0.32, 0.08] | -0.01 [-0.26, 0.26] | -0.00 [-0.26, 0.24] |
| Parent Value PE x Condition | -0.14 [-0.30, 0.03] | 0.16 [-0.05, 0.37] | 0.19 [-0.01, 0.39] |
| Friend Value PE x Condition | 0.14 [-0.02, 0.30] | -0.12 [-0.33, 0.07] | -0.14 [-0.33, 0.06] |

*Note.* Parameter estimates for the intercept, reward, and risk terms are not reported. ‘PE’ refers to pattern expression scores, obtained by using each individual subject’s parent and friend neural representations and a value-based neural signature. ‘Samp Specific’ refers to the type of signature used (Samp specific = sample-specific signature built using ridge regression). ‘2mm’ and ‘4mm’ refer to the degree of smoothing applied to the sample specific signature (the left-most column had no smoothing applied). Values in brackets represent 89% highest density credible intervals.

**Table 2.**
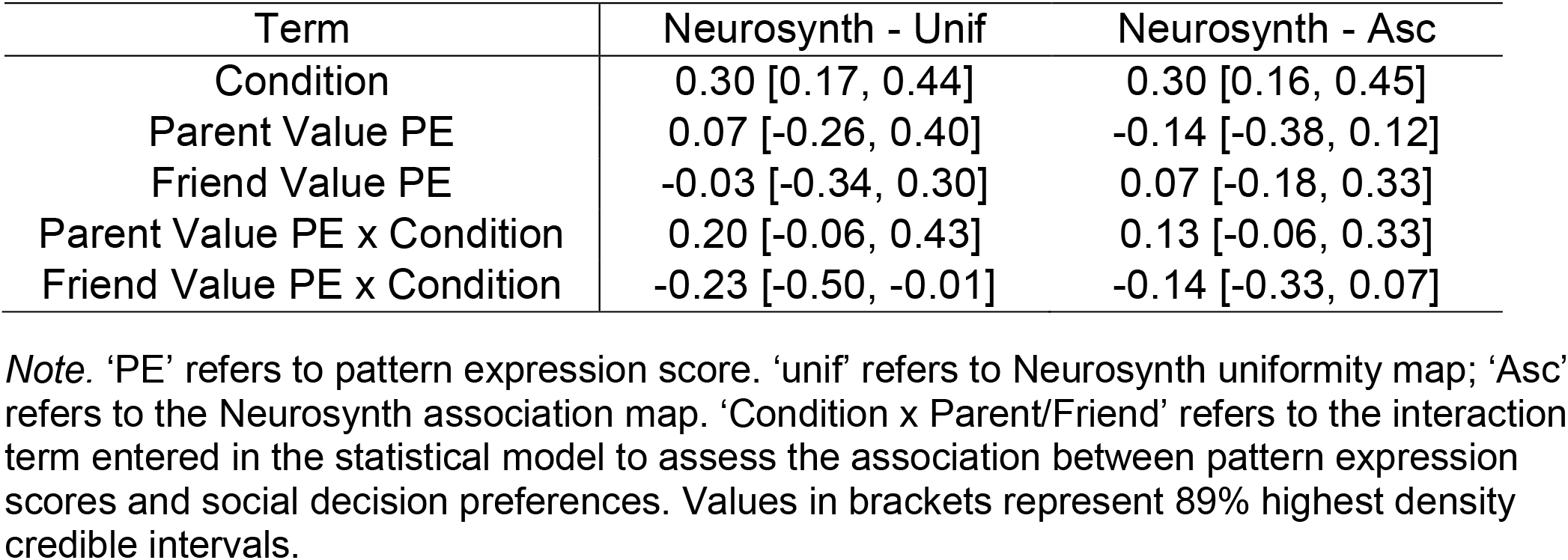
Predicting social decision preferences as a function of value-based representations using a Neurosynth meta-analytic neural signature.

**Figure 4.**
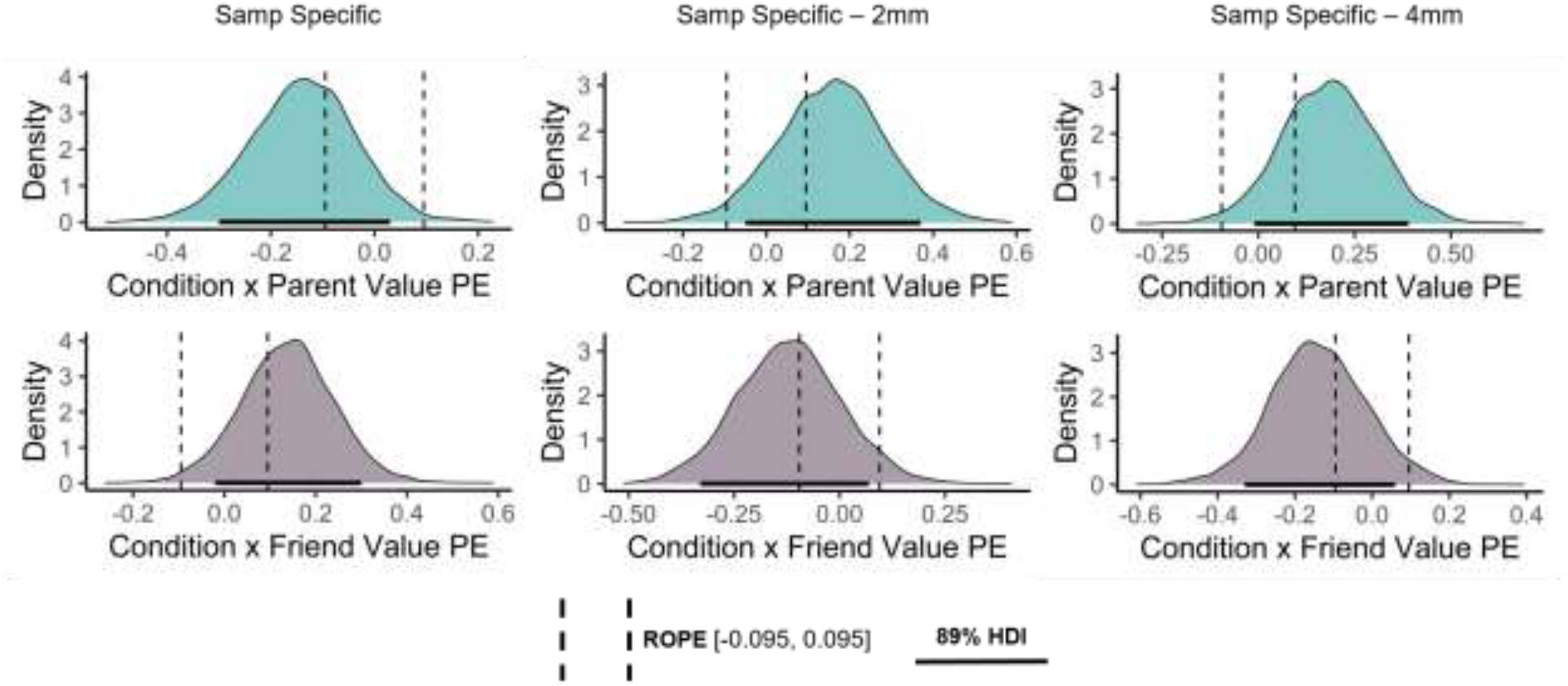
Posterior distribution plots for model interaction terms capturing the influence of value-based representations on social decision preferences (sample-specific neural signature). *Note*. ‘Samp Specific’ refers to the type of signature used (Samp specific = sample-specific signature built using ridge regression). ‘2mm’ and ‘4mm’ refer to the degree of smoothing applied to the sample specific signature (the left-most signature had no smoothing applied). ‘PE’ refers to pattern expression score. ‘Condition x Parent/Friend’ refers to the interaction term entered in the statistical model to assess the association between pattern expression scores and social decision preferences. ‘ROPE’ refers to Region of Practical Equivalence; ‘HDI’ refers to highest density credible intervals.

**Figure 5.**
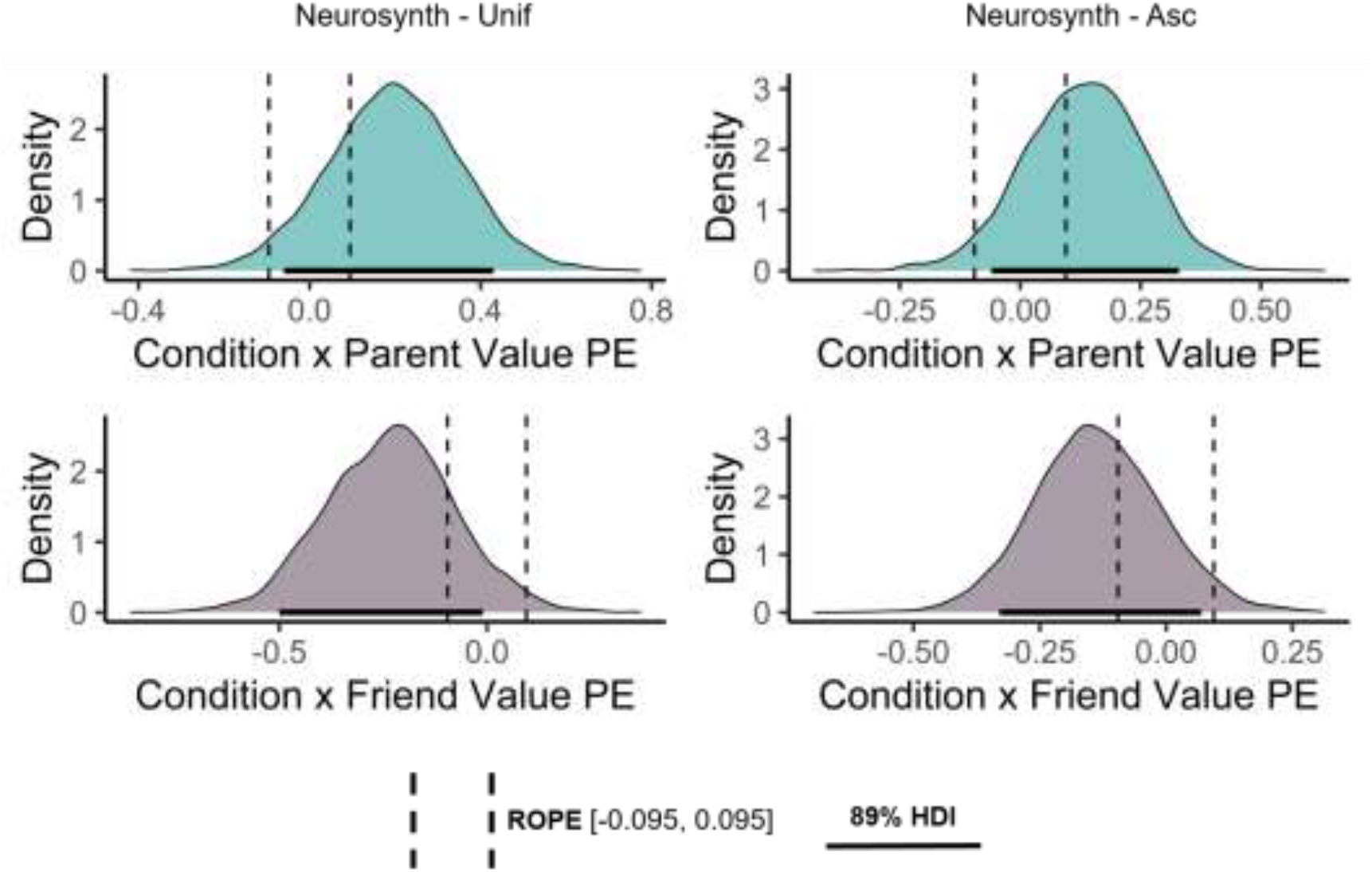
Posterior distribution plots for model interaction terms capturing the influence of value-based representations on social decision preferences (sample-specific neural signature). *Note*. ‘Unif’ refers to Neurosynth uniformity map; ‘Asc’ refers to the Neurosynth association map. ‘PE’ refers to pattern expression score. ‘PE’ refers to pattern expression score. ‘Condition x Parent/Friend’ refers to the interaction term entered in the statistical model to assess the association between pattern expression scores and social decision preferences. ‘ROPE’ refers to Region of Practical Equivalence; ‘HDI’ refers to highest density credible intervals.

To be consistent with our approach to analyzing paired differences, we conducted two similar *post-hoc* analyses. This involved re-running our hierarchical model (i) with V1 voxels masked out when computing pattern expression scores as well as (ii) computing pattern expression values in reward-related ROIs only. Overall, the results of these two *post-hoc* analyses are largely consistent with each other, as well as the initial planned analysis: a greater pattern expression score for a given individual was related with a stronger propensity to favor them on the modified CCT (see Supplemental Information, Supplementary Tables 1-2).

## Discussion

The current study sought to test what drives social decision preferences among close others. Cumulative evidence from multiple pattern expression analyses suggest that social decision preferences between two close others (i.e., one’s parent versus friend) are predicted by the extent to which the brain represents said close others in terms of value. These findings carry important implications about how representations of social agents drive social decision-making, as well as how representations are distributed across the brain.

### Underscoring the role of neural representations in social decision making

Excitingly, this study is among the first to examine how neural representations of others influence social decision behavior. By and large, the primary focus of most prior work has been on how individuals process and respond to various features of social decisions, often as applied to unfamiliar or distant others(e.g., the value of each decision alternative, the degree of risk involved, beliefs about a social partner’s resources or their attitudes) (e.g., Chang et al., 2011; Crockett et al., 2017; Fareri et al., 2015). By contrast, the present study focused on how individuals represent decision partners themselves (specifically, neural representations). This is noteworthy for a few reasons. First, examining social representations is a direct way of parsing the mechanisms that underlie social decision preferences. Representations of others are, theoretically, the lens through which we perceive and contextualize other’s behavior (Amodio, 2019; Tamir & Thornton, 2018). For example, representing one’s parent in terms of value could indicate a subjective sense of their relationship being fulfilling and of their basic needs (Tottenham, 2020). However, further work is needed to interrogate when said social preferences emerge – for example, they could theoretically stem from a sense of gratitude, a desire to maintain relationship strength or an entirely different motivational process. Second, and relatedly, the putative mechanistic influence that representations may have on social decision preferences are likely generalizable across contexts because of evidence that representations of others are theoretically stable and domain-general (Tamir & Thornton, 2018). For example, psychological theories of human development posit that initial representations of caregivers become internalized, remain stable across contexts and time, and inform how future representations of others are established (Bretheron, 1985, 1992). Such theories are strengthened by neuroscientific evidence showing that representational entities—ranging from objects to concepts—are enduring and stable across contexts (Ward et al., 2018; Lin & Thornton, 2021).

The present findings also have implications for understanding the role that neural computations of value play in social cognition and behavior (Zerubavel et al., 2015). Together with prior work, our results suggest that humans view others, at least in part, in terms of how they satisfy their own individual needs, which in turn motivates social behavior (Amodio, 2019; Tamir & Thornton, 2018). Our findings suggest that using a value-based neural architecture to construct representations of others is an efficient manner of determining the association between other social agents and one’s own goals. In other words, it suggests that conserved neural circuitry supports value computations across both social and non-social contexts (i.e., if our representations of a close other are aligned with value, it could suggest that our relationship with them is goal fulfilling). Future work may further unpack links between value-based representations and social behavior by attempting to explicitly formalize how value-based processes are integrated into representations of others. For instance, future work could examine how social experiences contribute to value-based encoding of others as a means to understand how such an encoding comes to be in the first place (e.g., Gonzalez & Chang, 2019).

### Representations are distributed across the brain

Another key takeaway from these findings is that they suggest the brain is not simply relying on two or three node circuits to perform low dimensional computations over decision-level inputs during social decision-making (e.g., computing subject value of a safe or risky option based on the degree of reward, uncertainty, etc.) (Gangopadhyay et al., 2021; Rilling & Sanfey, 2011). Our results are instead consistent with the notion that representations themselves are intrinsically high dimensional, given that they require storing and integrating a wealth of information in order to make real time predictions (Kriegeskorte & Douglas, 2018). This suggests that finely coded, multivariate information about others is leveraged to guide behavior during social decision making. Future work might seek to build on our findings and leverage more sophisticated techniques to learn more about the mechanistic details of high dimensional representations. For example, researchers might use neural networks to construct artificial representations of social agents that vary based on different facets of network architecture. This approach could be used to track how manipulating representations predicts simulated changes in social decision-making, and would have the added benefit of creating formalized models of how representational information is used in decision-making. Alternately, one could leverage animal models of social decision-making (Ben-Ami Bartal et al., 2011; Dal Monte et al., 2020) to better decode the specific computations by which representations guide behavior by decomposing value-based processes into its constituent components (e.g., White & Monosov, 2016) and examining how each component may map onto distinct neuronal populations that also encode representations of others.

### Limitations

Like all studies, the current one has limitations that warrant discussion. First, the lack of consistent evidence for a group-level value-based bias for parent or friend representations in either direction was somewhat surprising. One reason behind this may be due to the fact that parent and friend relationships are highly heterogeneous from person to person, enough so that group-level effects of this sort may be misleading or simply non-informative. Second, confidence in the present results would be strengthened by replication in larger and more diverse samples in light of ongoing debates about power in fMRI research (Marek, Tervo-Clemmens, et al., 2022). Third, future work might build on our findings regarding representations of value by looking at expression of other cognitive or affective processes (e.g., modeling representations of others in terms of semantic information, etc), as well as testing whether the modality of value matters (e.g., social versus monetary).

### Conclusion

Social decision-making in everyday life is nuanced and complex. The present study sought to better incorporate this complexity into the neuroscience of social decision-making by examining how representations of close others influence social decision making behavior. We found that value-based processes may influence social choice behavior in part via neural representations of close others.

## Supplementary Information

### Methods

#### Parent-Friend Nomination and Stimuli Collection

Upon signing up for the study, participants were informed that the study involved making hypothetical decisions on the behalf of a parent and close friend, and were asked to nominate one of each. Participants were not allowed to nominate current romantic partners or family members as “friends” in effort to avoid potential confounds. Afterwards, participants were asked to for the names they use to address their parent and friend, respectively, and for five ‘passport style’ headshots of each close other (each from a different angle). Images required neutral facial expressions, both eyes to be open, mouth shut, eyes locked straight ahead, and no head tilt (See Supplementary Figure 1 for an example). The experimenter first reviewed these requirements with participants via telephone and then sent them a PDF file with complete, detailed instructions (osf.io/muv2c). The experimenter assessed images for quality prior to the scan and asked participants for re-shoots if images did not comply with requirements.

#### Social Decision-making Paradigm

Consistent with our prior work (Guassi Moreira et al., 2018, 2020), we used a modified version of the computerized “hot” Columbia Card Task (CCT) to assess social decision-making preferences involving conflicting outcomes for parents and friends (Figner et al., 2009; van Duijvenvoorde et al., 2015). Participants completed two runs of the CCT, each consisting of 24 rounds. A single round is comprised of a series of iterative decisions, ranging between one and sixteen. During each round, participants were shown a set of sixteen overturned cards (i.e., collectively, a deck) and were told the objective of the task was to win points by iteratively turning over cards. Participants were informed that each card is associated with a gain or loss of points, and were made aware of a descriptive header above each deck that indicated the total number of loss in the deck (one or two), the point value of each loss card (-30 or -60), the point value of each gain card (10 or 20), and a running total meant to keep track of points earned so far on that deck. The configuration of loss cards, loss and gain values were crossed, yielding eight distinct deck types. Participants made choices regarding each deck type three times, hence 24 rounds.

#### Each round began with a score of zero points and all cards overturned

Participants were required to choose between turning over a card—a risky choice—or not turning over a card (‘passing’, a safe choice). If participants chose to turn over a card, the computer randomly selected a card and turned it over. Choosing to pass, by contrast, ended the round and participants could not gain or lose any additional points (akin to ‘cashing out’ at a casino). Each round lasted until the participant decided to pass or randomly flipped a loss card. Participants were informed the computer selected cards to flip at random. In reality, the first three risky choices for any deck were always rigged to flip a gain card to safeguard against participants losing too early and then feeling disproportionately discouraged from taking further risks. Participants completed four practice rounds to ensure proper understanding of the task. A trained experimenter did not allow them to proceed unless they demonstrated a clear understanding of the rules.

As previously mentioned, the CCT was modified to assess late adolescents’ social decision-making preferences between parents and friends. During one run of the task, participants were informed all points associated with gain cards would be awarded to their nominated parent, whereas any losses associated with each loss card would be incurred by their nominated friend. The opposite was true during the second run (gains solely benefit friend, losses solely incurred by parent). Critically, this manipulation models real-world trade-offs as participants were forced to make decisions that benefitted a close other at the potential expense of a second close other. The run order of the two conditions (Parent Gain-Friend Lose, Friend Gain-Parent Lose) was counterbalanced between subjects to ensure ordering of conditions did not affect decision behavior. There was never a trial in which only one close other is affected, ensuring there is always a potential cost for favoring one close other. Text describing the condition of the current run (‘Parent Gain | Friend Lose’ or ‘Friend Gain | Parent Lose’) was presented at the bottom of the screen on each trial as a reminder to participants. Outcomes were also clearly labeled to ensure participants understood what each close other may have won or lost following a given set of cards. The task was programmed and administered using the open-source, python-based PsychoPy software (Peirce, 2007). An experimenter remained present and unobtrusively monitor the participant during completion of the task in order to ensure participant focus and diligence.

#### Head Motion Statistics

Overall, head motion in this sample was low. For volumes exceeding the FD threshold, mean and maximum FD values were computed within each subject for each fMRI task. The means of these intra-subject metrics were used as a descriptive metric to reflect the overall head motion in the sample. The mean intra-subject average of *n* volumes exceeding the FD threshold on the parent-friend representation eliciting task was 0.484mm. The mean intra-subject average of the maximum FD value for this task was 0.620mm. Substantively, this means an ‘average subject’ is expected to move less than one volume above the FD threshold per run, and that their maximum FD value per run is expected to be ∼ 0.6mm. Only fifteen subjects exceeded the FD threshold during any run of the parent-friend representation eliciting task. For the coin flip task, the mean intra-subject average of *n* volumes exceeding the FD threshold was 0.467, and the mean intra-subject average of the maximum FD value was 0.637mm^1^. Twelve subjects exceeded the frame displacement threshold during any run of this task, whereas the rest did not.

#### Defining a Neural Signature of Value via Meta-Analysis (Neurosynth)

A second neural value signature was defined using meta-analytic maps from the online Neurosynth platform (Yarkoni et al., 2011). Neurosynth is an automated tool that extracts coordinates of brain activity from an actively maintained database of 14,371 studies (last updated July 2018), extracts high frequency terms occurring in the database’s studies, and uses this information to conduct a meta-analysis of activations for each term. Two images are computed for any given term: a uniformity image and association image. The uniformity map captures the degree of activity in the brain for a given term (comparable to how one would interpret results from a ‘standard’ whole-brain, univariate analysis). The association map is more selective, as it controls for base rates (e.g., quantifies how much more likely a given brain region is likely to be activated for a given term relative to studies that don’t include that term). More detailed on the platform can be accessed at neurosynth.org/faq. For this study, we used the meta-analytic map for the term ‘reward’. According to Neurosynth, 922 individual studies contributed to this term’s meta-analysis when our team downloaded the maps in late 2020. Both maps, uniformity and association, were used in analyses we report here.

#### Modeling Social Decision Preferences in a Hierarchical Bayesian Framework

Decisions on the i-th trial from the j-th participant on the modified CCT were modeled as being distributed Bernoulli.

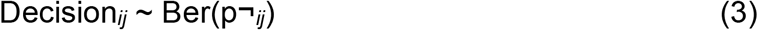

The Bernoulli distribution is frequently used to model binary outcomes, and takes a single parameter (p) describing the probability of ‘success’. Here, pij represents the probability of the j-th participant making a risky decision (i.e., turning over a card) on the -th trial. The log odds of these probabilities were further modeled as a linear combination of trial-level variables; an intercept (b0j), the experimental condition (b1j; 1 = Parent Gain-Friend Lose, 0 = Friend Gain-Parent Lose), return (b2j), and risk (b3j).

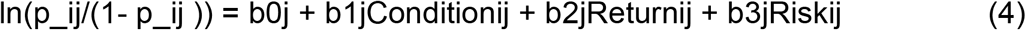

Critically, b1j is the key parameter of interest, as it encodes social decision preferences. A value equal to zero indicates no preference, a positive value indicates a parent-over-friend preference, and a negative value indicates a friend-over-friend preference. Return represents the expected value associated with flipping over a card on the i-th trial; risk represents the variance in the outcome distribution for the i-th trial. Modeling both is a common practice that helps statistically adjust for decision-level features as well as provides a ‘sanity check’ that participants are completing the task correctly (van Duijvenvoorde et al., 2015) Coefficients represent expected changes in logit units – that is, a one unit increase in any predictor will be associated with an expected change in the log odds of a risky decision equivalent to b (referring to a generic coefficient). Logit units can be converted to an odds ratio (i.e., the expected change in the odds) by exponentiating a given coefficient (i.e., exp(b)).

Notably, coefficients associated with these trial-level variables can be decomposed into population-level (γ) and group-level (u) parameters, loosely analogous to the concept of fixed and random effects. Between-subject predictors were included in the model, as moderators of the effect of condition (social decision preferences). This means the slopes associated with the intercept (b0j) and condition (b1j) were parameterized as equations (5-6).

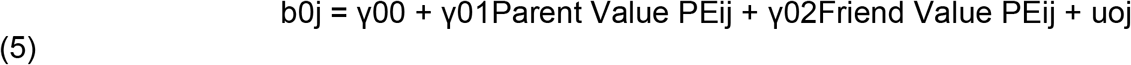

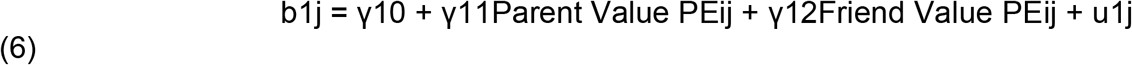

Here, Parent Value PE refers to value-based pattern expression scores for the parent representation, and Friend Value PE refers to same quantity but with friend representations. Between-subjects predictors were not added to the slopes for return (b2j) and risk (b3j), as notated in equations (7-8).

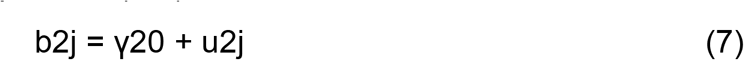

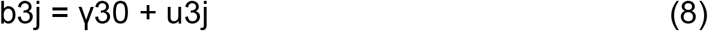

Hierarchical models were fit using the default sampling procedures in the brms R package (no thinning, 4 chains, 4000 samples per chains, 2000 discarded warm-up samples).

When selecting priors for the hierarchical model, we hoped to achieve two goals: (i) avoid adding substantial bias to the analysis (e.g., the prior suggests an effect that is not actually present), (ii) achieve principled regularization of model parameters. While regularization is traditionally used in the context of predictive modeling (e.g., Hine & Usynin, 2005; Stiglic et al., 2015; Xiao et al., 2018; Yao & Yang, 2016), regularized approaches can also aid inference by minimizing the influence of noise in parameter estimation and therefore enhancing parameter generalizability (Efron & Morris, 1975, 1977; James & Stein, 1992). This left us to consider two types of prior distributions: non-informative and weakly informative. Non-informative priors assume all parameter values are equally likely, whereas weakly informative priors are modestly confident about which parameter values are more likely than not. These categories are in contrast to informative priors which encode highly specific beliefs about model parameters and, in our case here, carried greater potential to bias the analysis. We ultimately selected weakly informative priors because the non-informative prior is too diffuse to regularize parameter estimates and is more likely to lead to inappropriately high posterior mass around extreme, highly implausible parameter value.

Thus, all fixed effects received a standard normal prior (N(0,1)). The normal distribution was selected because we did not have reason to suspect asymmetry in the parameter space and did not have a reason to believe that fatter tails (e.g., in a t-distribution) were necessary given the logistic regression model. The location and scale (mean, standard deviation) parameters of 0 and 1 were selected because (i) zero corresponds with the null value and regularization typically occurs by biasing coefficients to a null value, and (ii) a standard deviation of 1—in logistic regression— would virtually cover the entire range of plausible parameter estimates (effects greater than |3| in logistic regression correspond to enormous effect sizes on an odds scale, certainly larger than would be expected in behavioral science research). The random effects from the model were drawn from a student’s distribution (t(3, 0, 2.5)). This t-distribution was used at the recommendation of the brms developer, who notes that group-level effects often require distributions with fatter tails.

#### Deriving a Sample-Specific Neural Signature of Value

We started to create a sample-specific neural signature of value by concatenating single trial activations from all subjects in a *time* by *voxel* matrix, where each row represented a single trial from the *i*-th subject, and each column represented a specific voxel. The matrix was reduced using PCA, following the precedent established by other similar studies (Chang et al., 2015; Krishnan et al., 2016; Wager et al., 2013), resulting in 1500 components that comprised 90% of the explained variance from the original matrix. LASSO (Least Absolute Shrinkage and Selection Operator) and ridge regression were used to predict each trial’s monetary value from BOLD activity, indexed by the set of principle components.

Ridge and LASSO regression were selected for several reasons. Foremost, the nature of the data demanded an analytic method that could handle continuous outcomes. Second, both methods use penalized estimators, which have the effect of regularizing parameter estimates (i.e., biasing them towards, or to, zero in order to reduce variability in sample-to-sample estimates). As previously mentioned, this helps enhance the generalizability of parameter estimates and safeguards against overfitting. Third, they are broadly consistent with existing, similar studies (Chang et al., 2015; Krishnan et al., 2016; Wager et al., 2013). Last, these models can handle highly parameterized models without encountering estimation problems (e.g., parameter estimate instability).

We used 10-fold cross-validation to determine the best penalty for both ridge and LASSO models. After obtaining the ideal tuning parameter, a model using each type of estimator was fit to the complete dataset predicting monetary value on each trial from principle components of brain activity (indexed via the BOLD signal). Evaluating both models using the *R*^*2*^ metric of model fit, we found that the ridge regression model fit the data better than LASSO. We backtransformed the weights of each principal component into the original voxel space, and thresholded the weights at zero^2^, creating the final neural signature map. Visually inspecting the final ridge regression-based neural signature revealed some degree of sparsity among the voxel weights. Realizing this could be a potential signal-to-noise issue, we created two additional versions of the map by spatially smoothing the weights using a 2mm and 4mm Gaussian kernel (fwhm). Subsequent analyses using the sample-specific signature report results using all three of these models. The unsmoothed signature is depicted in Figure 2 in the main document.

To further check whether this sample-specific map was properly indicative of reward, we correlated the weights in our map with meta-analytic maps of reward and of four other terms: language, pain, working memory and social (all obtained via Neurosynth). The correlations between the neural signature of value and meta-analytic maps from unrelated constructs (language, pain, working memory, social) were low in magnitude (non-smoothed signature *r*s = -0.068 – 0.001; 2mm-smoothed signature *r*s = -0.063 – 0.018; 4mm-smoothed signature *r*s = -0.052 – 0.019), whereas the correlation between the reward map and the signature was higher (non-smooth signature *r* = 0.306; 2mm-smoothed signature *r* = 0.356; 4mm-smoothed signature *r* = 0.505). This provides discriminant and converging evidence that the signature measures what it is intended to. Further, that the correlation with the meta-analytic map of reward was not very high (e.g., >.7), suggests our signature could be capturing unique or distinct facets of valuation (i.e., it is not redundant with the meta-analytic map). Visual inspection of the signature shows regions canonically associated with reward (e.g., striatum, medial prefrontal cortex) are present in the anticipated direction, further suggesting the signature is at least measuring the intended psychological process.

### Results

#### Manipulation Checks

In terms of behavioral social decision preferences, we used the same modeling framework as described in the main texts and this supplement. We observed evidence for a mean-level parent-over-friend social decision preference (posterior mean of social decision preference parameter: 0.30, 89% CI = [0.16, 0.44]).

Imaging results obtained using a mixed effects model (FSL’s FLAME1) and subsequently cluster-corrected using random field theory (Family-wise-error < .05, cluster defining threshold Z > 3.1) show robust activation in the ventral striatum (k = 1570, L: x = -14 y = 6 z = -10, Z = 5.516 ; R: x = 16 y = 4 z = -12, Z = 5.517) as well as the medial prefrontal cortex (k = 539, x = -4 y = 56 z = 2, Z = 4.500) for the Win > Loss contrast, indicating a successful replication of prior work (Haber & Knutson, 2009; Knutson et al., 2001; Supplementary Figure 2). This suggests the desired psychological state was evoked during the task rendering the data suitable for attempting to define a sample-specific neural signature of value.

#### Post-Hoc Analyses for Paired Differences in Value-Based Pattern Expression of Neural Representations

The first post-hoc analysis involved excluding primary visual cortex (V1) from the neural signature, under the reasoning that visual processes are unlikely to reflect meaningful information about valuation (Kragel et al., 2019). However, re-running the pattern expression analysis with a custom neural signature that excluded V1 voxels did not meaningfully change the results (Samp specific: posterior mean, (SD): d = -0.07 (0.14), 89% CI: [-0.29, 0.17]) (Samp specific – 2mm: posterior mean, (SD): d = -0.04 (0.14), 89% CI: [-0.27, 0.20]) (Samp specific – 4mm: posterior mean, (SD): d = - 0.04 (0.14), 89% CI: [-0.26, 0.19]). The second post-hoc analysis masked brain regions in the custom signature thought to be central to valuation, the ventral striatum (VS) and medial prefrontal cortex (mPFC) (Dabney et al., 2020; Haber & Knutson, 2009; Kutlu et al., 2021; Lopez-Persem et al., 2020; Vlaev et al., 2011). However, this analysis again yielded equivocal evidence for a value-based signature bias in either direction (Samp specific: posterior mean, (SD): d = 0.07 (0.14), 89% CI: [-0.16, 0.30]) (Samp specific – 2mm: posterior mean, (SD): d = 0.02 (0.14), 89% CI: [-0.21, 0.25]) (Samp specific – 4mm: posterior mean, (SD): d = 0.02 (0.14), 89% CI: [-0.25, 0.21]).

*Supplementary Figure 1.* Sample example stimuli.

FIGURE OMITTED DUE TO BIORXIV POLICY OVER IDENTIFIABLE INFORMATION

*Note*. These images were taken from the sample PDF instruction file. These images are not actual subject data.

**Supplementary Figure 2.**
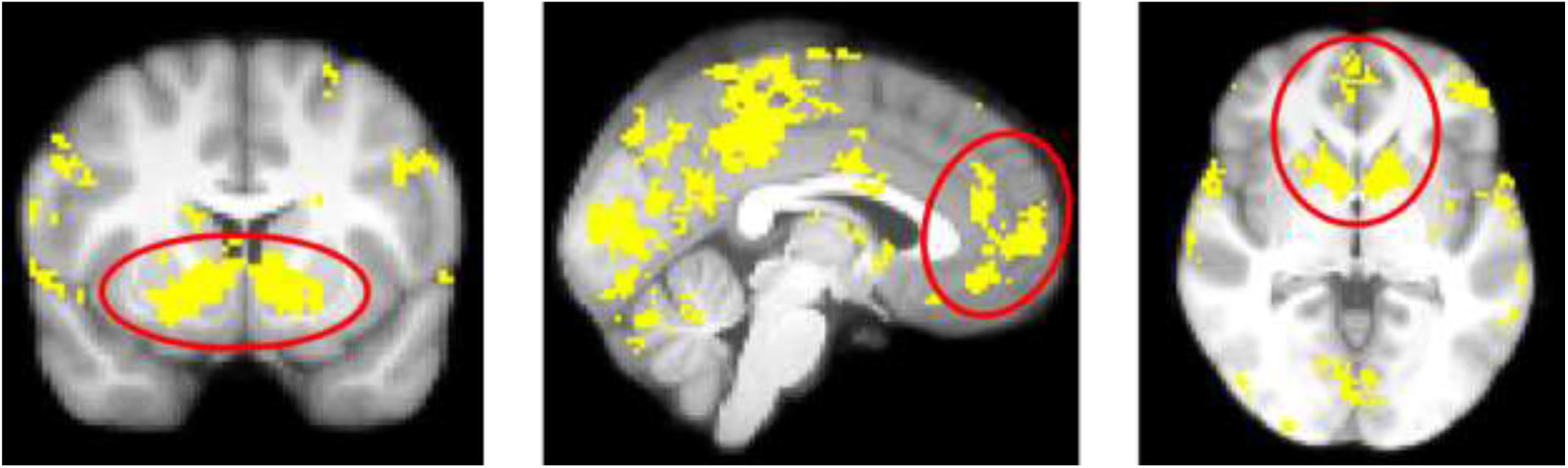
Results of Win>Loss contrast during the coin flip task (value) *Note*. Winning, relative to losing, on the coin flip task evoked robust activity in the ventral striatum and medial prefrontal cortex (circled in red). Cluster corrected (Family-Wise-Error of *p* < .05) using FLS’s FLAME1 (Cluster Defining Threshold of Z > 3.1).

**Supplementary Figure 3.**
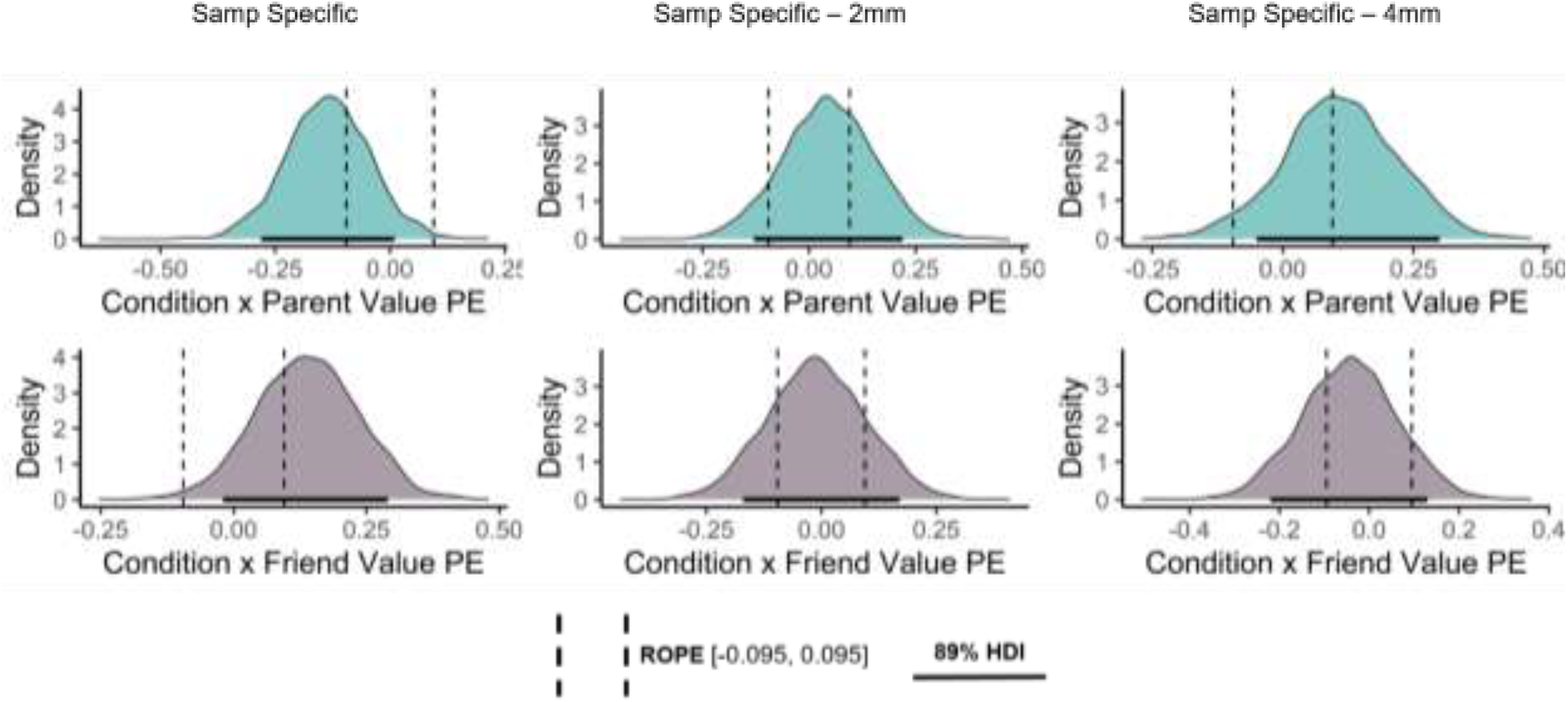
Posterior distribution plots for model interaction terms capturing the influence of value-based representations on social decision preferences (sample-specific neural signature, excluding V1 – post-hoc analysis). *Note*. ‘Samp Specific’ refers to the type of signature used (Samp specific = sample-specific signature built using ridge regression). ‘2mm’ and ‘4mm’ refer to the degree of smoothing applied to the sample specific signature (the left-most signature had no smoothing applied). ‘PE’ refers to pattern expression score. ‘Condition x Parent/Friend’ refers to the interaction term entered in the statistical model to assess the association between pattern expression scores and social decision preferences. ‘ROPE’ refers to Region of Practical Equivalence; ‘HDI’ refers to highest density credible intervals.

**Supplementary Figure 4.**
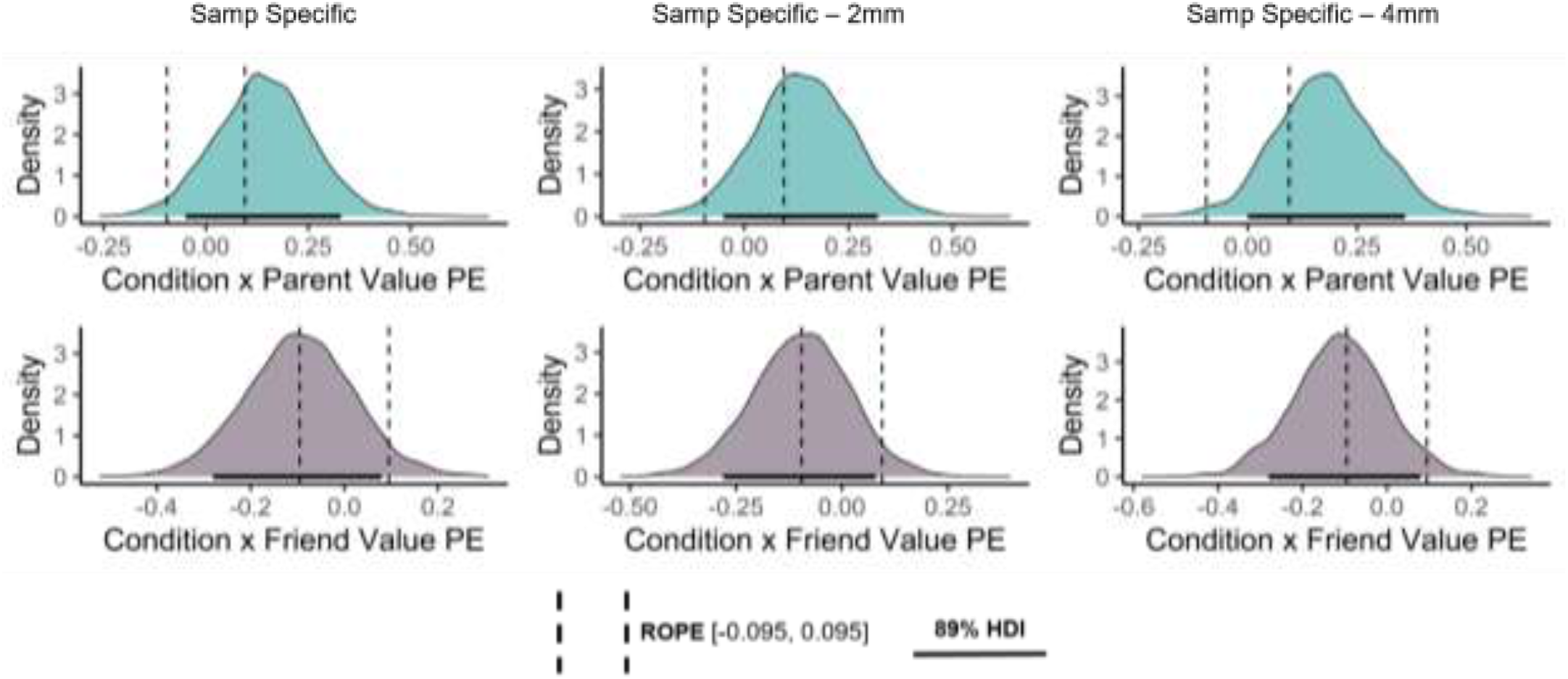
Posterior distribution plots for model interaction terms capturing the influence of value-based representations on social decision preferences (sample-specific neural signature, including only VS, mPFC – post-hoc analysis). *Note*. ‘Samp Specific’ refers to the type of signature used (Samp specific = sample-specific signature built using ridge regression). ‘2mm’ and ‘4mm’ refer to the degree of smoothing applied to the sample specific signature (the left-most signature had no smoothing applied). ‘PE’ refers to pattern expression score. ‘Condition x Parent/Friend’ refers to the interaction term entered in the statistical model to assess the association between pattern expression scores and social decision preferences. ‘ROPE’ refers to Region of Practical Equivalence; ‘HDI’ refers to highest density credible intervals.

**Supplementary Table 1.**
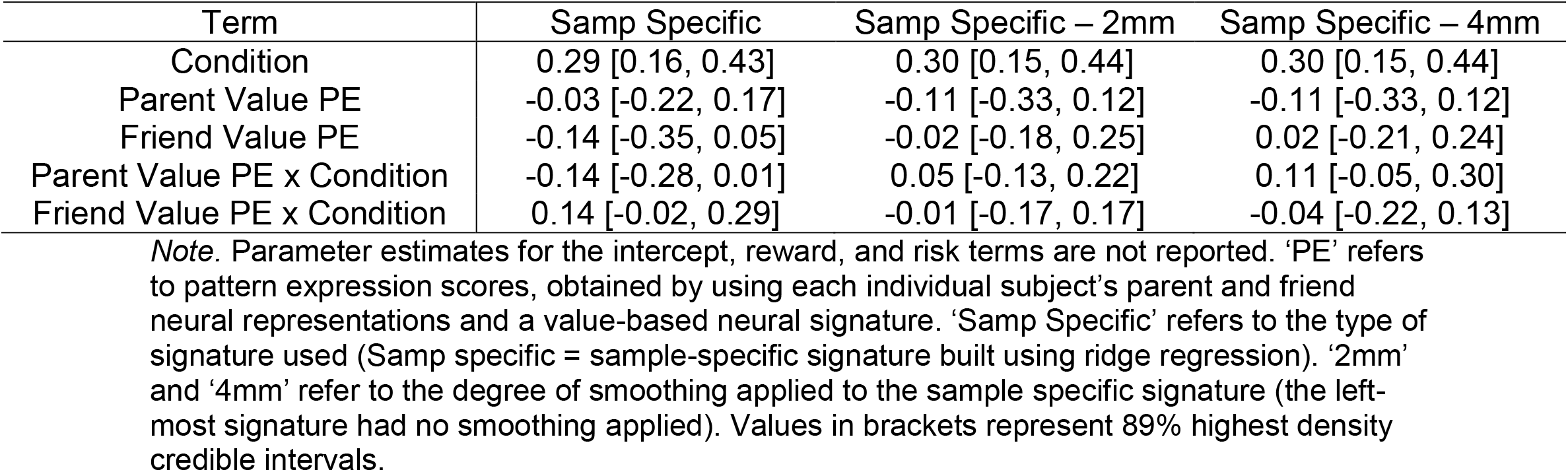
Predicting social decision preferences as a function of value-based representations using a sample-specific neural signature, excluding primary visual cortex (V1 – post-hoc analysis)

**Supplementary Table 2.** Predicting social decision preferences as a function of valuebased representations using a sample-specific neural signature, including only reward regions (VS, mPFC – post-hoc analysis).

| Term | Samp Specific | Samp Specific – 2mm | Samp Specific – 4mm |
| --- | --- | --- | --- |
| Condition | 0.30 [0.16, 0.44] | 0.30 [0.16, 0.44] | 0.30 [0.16, 0.44] |
| Parent Value PE | -0.03 [-0.27, 0.21] | -0.00 [-0.23, 0.24] | -0.03 [-0.28, 0.20] |
| Friend Value PE | -0.04 [-0.27, 0.19] | -0.05 [-0.28, 0.18] | -0.02 [-0.25, 0.23] |
| Parent Value PE x Condition | 0.18 [0.00, 0.36] | 0.14 [-0.05, 0.32] | 0.14 [-0.05, 0.33] |
| Friend Value PE x Condition | -0.11 [-0.28, 0.08] | -0.10 [-0.28, 0.08] | -0.09 [-0.28, 0.08] |

*Note.* Parameter estimates for the intercept, reward, and risk terms are not reported. ‘PE’ refers to pattern expression scores, obtained by using each individual subject’s parent and friend neural representations and a value-based neural signature. ‘Samp Specific’ refers to the type of signature used (Samp specific = sample-specific signature built using ridge regression). ‘2mm’ and ‘4mm’ refer to the degree of smoothing applied to the sample specific signature (the left-most signature had no smoothing applied). Values in brackets represent 89% highest density credible intervals. ‘VS’ refers to ventral striatum, mPFC refers to medial prefrontal cortex.

## Footnotes

1 Based on participant feedback received during two pilot scans, participants were asked to provide the labels they use to address each close other (e.g., ‘Mom’ or ‘Dad’ for a parent).

2 All other GLM specifications (e.g., slice timing correction via temporal derivatives, regressor convolution, etc.) for the LSS analysis were identical to those used in the parent-friend representation GLMs.

1 This estimate excludes the aforementioned outlying participant who averaged 20+ volumes exceeding the FD threshold.

2 It was difficult to conceptualize what a negative association between brain responses and coin flip task values could indicate.

## References

Blakemore, S.-J., & Mills, K. L. (2014). Is adolescence a sensitive period for sociocultural processing? Annual Review of Psychology, 65, 187–207. https://doi.org/10.1146/annurev-psych-010213-115202

Braams, B. R., & Crone, E. A. (2016). Peers and parents: A comparison between neural activation when winning for friends and mothers in adolescence. Social Cognitive and Affective Neuroscience, [Epub ahead of print].

Chang, L. J., Gianaros, P. J., Manuck, S. B., & Krishnan, A. (2015). A sensitive and specific neural signature for picture-induced negative affect. PLOS Biology, 13(6), 1–28. https://doi.org/10.1371/journal.pbio.1002180

Chang, L. J., Smith, A., Dufwenberg, M., & Sanfey, A. G. (2011). Triangulating the neural, psychological, and economic bases of guilt aversion. Neuron, 70(3), 560–572.

Charest, I., Kievit, R. A., Schmitz, T. W., Deca, D., & Kriegeskorte, N. (2014). Unique semantic space in the brain of each beholder predicts perceived similarity. Proceedings of the National Academy of Sciences of the United States of America, 111(40), 14565–14570. https://doi.org/10.1073/pnas.1402594111

Chavez, R. S., Heatherton, T. F., & Wagner, D. D. (2017). Neural population decoding reveals the intrinsic positivity of the self. Cerebral Cortex, 27(11), 5222–5229. https://doi.org/10.1093/cercor/bhw302

Chen, G., Taylor, P. A., & Cox, R. W. (2017). Is the statistic value all we should care about in neuroimaging? NeuroImage, 147, 952–959. https://doi.org/10.1101/064212

Clark, A., & Toribio, J. (1994). Doing without representing? Synthese, 101(3), 401–431.

Cosme, D., Zeithamova, D., Stice, E., & Berkman, E. T. (2019). Multivariate neural signatures for health neuroscience: Assessing spontaneous regulation during food choice. Social Cognitive and Affective Neuroscience, in press.

Cremers, H. R., Wager, T. D., & Yarkoni, T. (2017). The relation between statistical power and inference in fMRI. PLoS ONE, 12(11), e0184923.

Crockett, M. J., Kurth-Nelson, Z., Siegel, J. Z., Dayan, P., & Dolan, R. J. (2014). Harm to others outweighs harm to self in moral decision making. Proceedings of the National Academy of Sciences, 111(48), 17320–17325. https://doi.org/10.1073/pnas.1424572112

Crockett, M. J., Siegel, J. Z., Kurth-Nelson, Z., Dayan, P., & Dolan, R. J. (2017). Moral transgressions corrupt neural representations of value. Nature Neuroscience, May, 1–10. https://doi.org/10.1038/nn.4557

DeCharms, R. C., & Zador, A. (2000). Neuro representation and the cortical code. Annual Review of Neuroscience, 23, 613–647.

Doré, B. P., Weber, J., & Ochsner, K. N. (2017). Neural predictors of decisions to cognitively control emotion. Journal of Neuroscience, 37(10), 2580–2588. https://doi.org/10.1523/JNEUROSCI.2526-16.2016

Etzel, J. A., Valchev, N., & Keysers, C. (2011). NeuroImage The impact of certain methodological choices on multivariate analysis of fMRI data with support vector machines. NeuroImage, 54(2), 1159–1167. https://doi.org/10.1016/j.neuroimage.2010.08.050

Fareri, D. S., Chang, L. J., & Delgado, M. R. (2015). Computational substrates of social value in interpersonal collaboration. Journal of Neuroscience, 35(21), 8170–8180.

Fareri, D. S., Smith, D. V., & Delgado, M. R. (2020). The influence of relationship closeness on default-mode network connectivity during social interactions. Social Cognitive and Affective Neuroscience, 15(3), 261–271.

Fareri, D. S., Stasiak, J. E., & Sokol-Hessner, P. (2022). Choosing for others changes dissociable computational mechanisms underpinning risky decision-making. PsyArXiv.

Feldmanhall, O., & Chang, L. J. (2018). Social Learning: Emotions Aid in Optimizing Goal-Directed Social Behavior. In R. Morris, A. Bornstein, & A. Shenhav (Eds.), Goal-Directed Decision-Making: Computations and Circuits (pp. 309–330). Academic Press. https://doi.org/10.1016/B978-0-12-812098-9.00014-0

Figner, B., Mackinlay, R. J., Wilkening, F., & Weber, E. U. (2009). Affective and deliberative processes in risky choice: age differences in risk taking in the Columbia Card Task. Journal of Experimental Psychology. Learning, Memory, and Cognition, 35(3), 709–730. https://doi.org/10.1037/a0014983

Gee, D. G., Gabard-Durnam, L., Telzer, E. H., Humphreys, K. L., Goff, B., Shapiro, M., Flannery, J., Lumian, D. S., Fareri, D. S., Caldera, C., & Tottenham, N. (2014). Maternal buffering of human amygdala-prefrontal circuitry during childhood but not during adolescence. Psychological Science, 25(11), 2067–2078. https://doi.org/10.1177/0956797614550878

Gonzalez, B., & Chang, L. J. (2019). Computational models of mentalizing. PsyArXiv, 1–19.

Guassi Moreira, J. F., Tashjian, S. M., Galvan, A., & Silvers, J. A. (2021). Computational and motivational mechanisms of human social decision making involving close others. Journal of Experimental Social Psychology, 93, 1–10.

Guassi Moreira, J. F., Tashjian, S. M., Galván, A., & Silvers, J. A. (2018). Parents versus peers: Assessing the impact of social agents on decision making in young adults. Psychological Science, 29(9), 1526–1539. https://doi.org/10.1177/0956797618778497

Guassi Moreira, J. F., Tashjian, S. M., Galván, A., & Silvers, J. A. (2020). Is social decision making for close others consistent across domains and within individuals? Journal of Experimental Psychology: General, 149(8), 1509–1526.

Guthrie, T., Benadjaoud, Y. Y., & Chavez, R. S. (2022). Social relationship strength modulates the similarity of brain-to-brain representations of group members. Social Cognitive and Affective Neuroscience, 32(11), 2469–2477.

Haber, S. N., & Knutson, B. (2009). The Reward Circuit: Linking Primate Anatomy and Human Imaging. Neuropsychopharmacology, 35(10), 4–26. https://doi.org/10.1038/npp.2009.129

Hackel, L. M., Zaki, J., & Van Bavel, J. J. (2017). Social identity shapes social valuation: evidence from prosocial behavior and vicarious reward. Social Cognitive and Affective Neuroscience, 12(8), 1219–1228. https://doi.org/10.1093/scan/nsx045

Hong, Y., Yoo, Y., Han, J., Wager, T. D., & Woo, C.-W. (2019). False-positive neuroimaging: Undisclosed flexibility in testing spatial hypotheses allows presenting anything as a replicated finding. NeuroImage, 195, 384–395.

Johnson, N. D., & Mislin, A. A. (2011). Trust games: A meta-analysis. Journal of Economic Psychology, 32(5), 865–889. https://doi.org/10.1016/j.joep.2011.05.007

Knutson, B., Fong, G. W., Adams, C. M., Varner, J. L., & Hommer, D. (2001). Dissociation of reward anticipation and outcome with event-related fMRI. NeuroReport, 12(17), 3683–3687. https://doi.org/10.1097/00001756-200112040-00016

Kruschke, J. K. (2011). Bayesian assessment of null values via parameter estimation and model comparison. Perspectives on Psychological Science, 6(3), 299–312.

Kruschke, J. K. (2013). Bayesian estimation supersedes the t-test. Journal of Experimental Psychology: General, 142(2), 573–603.

Lamba, A., Frank, M. J., & FeldmanHall, O. (2020). Anxiety impedes adaptive social learning under uncertainty. Psychological Science. https://doi.org/10.1177/0956797620910993

Makowski, D., Ben-Shachar, M. S., & Ludecke, D. (2019). bayestestR: Describing effects and their uncertainty, existence and significance within the bayesian framework. Journal of Open Source Software, 4(40), 1541. https://doi.org/10.21105/joss.01541

McElreath, R. (2018). Statistical rethinking: A bayesian course with examples in R and Stan (1st ed.). Chapman and Hall/CRC. https://www.taylorfrancis.com/books/mono/10.1201/9781315372495/statistical-rethinking-richard-mcelreath

Misaki, M., Luh, W.-M., & Bandettini, P. A. (2014). The effect of spatial smoothing on fMRI decoding of columnar-level organization with linear support vector machine. Journal of Neuroscience Methods, 212(2), 355–361. https://doi.org/10.1016/j.jneumeth.2012.11.004.The

Morgan, A. (2014). Representations gone mental. Synthese, 191(2), 213–244.

Mumford, J.A. (2012). A power calculcation guide for fMRI studies. Social Cognitive and Affective Neuroscience, 7(6), 738–742.

Mumford, Jeanette A., Davis, T., & Poldrack, R. A. (2014). The impact of study design on pattern estimation for single-trial multivariate pattern analysis. NeuroImage, 103, 130–138. https://doi.org/10.1016/j.neuroimage.2014.09.026

Mumford, Jeanette A., Turner, B. O., Ashby, F. G., & Poldrack, R. A. (2012). Deconvolving BOLD activation in event-related designs for multivoxel pattern classification analyses. NeuroImage, 59(3), 2636–2643. https://doi.org/10.1016/j.neuroimage.2011.08.076

Ong, D. C., Zaki, J., & Gruber, J. (2017). Increased cooperative behavior across remitted bipolar I disorder and major depression: Insights utilizing a behavioral economic trust game. Journal of Abnormal Psychology, 126(1).

Parkinson, C., Kleinbaum, A. M., & Wheatley, T. (2017). Spontaneous neural encoding of social network position. Nature Human Behaviour, 1(5), 1–7. https://doi.org/10.1038/s41562-017-0072

Poldrack, R. A. (2021). The physics of representation. Synthese, 199, 1307–1325.

Poldrack, R. A., Baker, C. I., Durnez, J., Gorgolewski, K. J., Matthews, P. M., Munafò, M., Nichols, T. E., Poline, J.-B., Vul, E., & Yarkoni, T. (2017). Scanning the horizon: Towards transparent and reproducible neuroimaging research. Nature Reviews Neuroscience, 18, 115–126. https://doi.org/http://dx.doi.org/10.1101/059188

Powers, K. E., Yaffe, G., Hartley, C. A., Davidow, J. Y., Kober, H., & Somerville, L. H. (2017). Consequences for peers differentially bias computations about risk across development. Journal of Experimental Psychology: General, Online ahead of print. https://doi.org/10.1037/xge0000389

Siegel, J. S., Power, J. D., Dubis, J. W., Vogel, A. C., Church, J. A., Schlaggar, B. L., & Petersen, S. E. (2014). Statistical improvements in functional magnetic resonance imaging analyses produced by censoring high-motion data points. Human Brain Mapping, 35(5), 1981–1996. https://doi.org/10.1002/hbm.22307.Statistical

Steinberg, L., & Morris, A. S. (2001). Adolescent development. Annual Review of Psychology, 52, 83–110. https://doi.org/10.1146/annurev.psych.52.1.83

Tamir, D. I., & Thornton, M. A. (2018). Modeling the predictive social mind. Trends in Cognitive Sciences, 22(3), 201–212.

Taylor, M. J., Arsalidou, M., Bayless, S. J., Morris, D., Evans, J. W., & Barbeau, E. J. (2009). Neural correlates of personally familiar faces: Parents, partner and own faces. Human Brain Mapping, 30(7), 2008–2020. https://doi.org/10.1002/hbm.20646

van de Groep, S., Sweijen, S. W., de Water, E., & Crone, E. A. (2022). Temporal discounting for self and friend in adolescence: a fMRI study. PsyArXiv.

van Duijvenvoorde, A. C. K., Huizenga, H. M., Somerville, L. H., Delgado, M. R., Powers, A., Weeda, W. D., Casey, B. J., Weber, E. U., & Figner, B. (2015). Neural correlates of expected risks and returns in risky choice across development. The Journal of Neuroscience, 35(4), 1549–1560. https://doi.org/10.1523/JNEUROSCI.1924-14.2015

White, J. K., & Monosov, I. E. (2016). Neurons in the primate dorsal striatum signal the uncertainty of object-reward associations. Nature Communications, 7, 1–8. https://doi.org/10.1038/ncomms12735

Yarkoni, T., Poldrack, R. A., Nichols, T. E., Van Essen, D. C., & Wager, T. D. (2011). Large-scale automated synthesis of human functional neuroimaging data. Nature Methods, 8(8), 665–670.

Zerubavel, N., Bearman, P. S., Weber, J., & Ochsner, K. N. (2015). Neural mechanisms tracking popularity in real-world social networks. Proceedings of the National Academy of Sciences, 112(49), 15072–15077. https://doi.org/10.1073/pnas.1511477112

